# splicekit: a comprehensive toolkit for splicing analysis from short-read RNA-seq

**DOI:** 10.1101/2023.05.25.542256

**Authors:** Gregor Rot, Arne Wehling, Roland Schmucki, Nikolaos Berntenis, Jitao David Zhang, Martin Ebeling

## Abstract

**Motivation:** Analysis of alternative splicing using short-read RNA-seq data is a complex process that involves several steps: alignment of reads to the reference genome, identification of alternatively spliced features, motif discovery, analysis of RNA-protein binding near donor and acceptor splice sites, and exploratory data visualization.

**Results:** We introduce *splicekit*, a python package that provides a comprehensive set of tools for conducting splicing analysis.

**Availability and implementation:** https://github.com/bedapub/splicekit and over PyPI.

## 1 Introduction

Alternative splicing of RNA is a fundamental biological process that is critical for the generation of protein diversity. Dysregulation of splicing has been implicated in many human diseases such as cancer and neurological disorders (Scotti et al., 2016). Recent advances in splicing modulation using compounds, i.e. small molecules (Schneider-Poetsch et al., 2021), such as Risdiplam for the treatment of spinal muscular atrophy (Ratni et al., 2018), have renewed interest in developing new therapies targeting splicing.

Splicing analysis using short-read RNA-seq data is a multifaceted process that involves several steps and requires the integration of diverse software tools. For the analysis of differential splicing events, the community can potentially benefit from a comprehensive and efficient analysis toolbox.

To address this need, we introduce *splicekit*, a Python package that provides and integrates a set of existing and novel splicing analysis tools (Figure 1A). It offers functionalities to identify differentially expressed features (junctions, exons and genes), cluster samples, perform motif analysis to elucidate potential regulatory patterns, visualize changes of junction versus those of genes, and to identify RNA-protein binding in the vicinity of regulated features.

**Figure 1.**
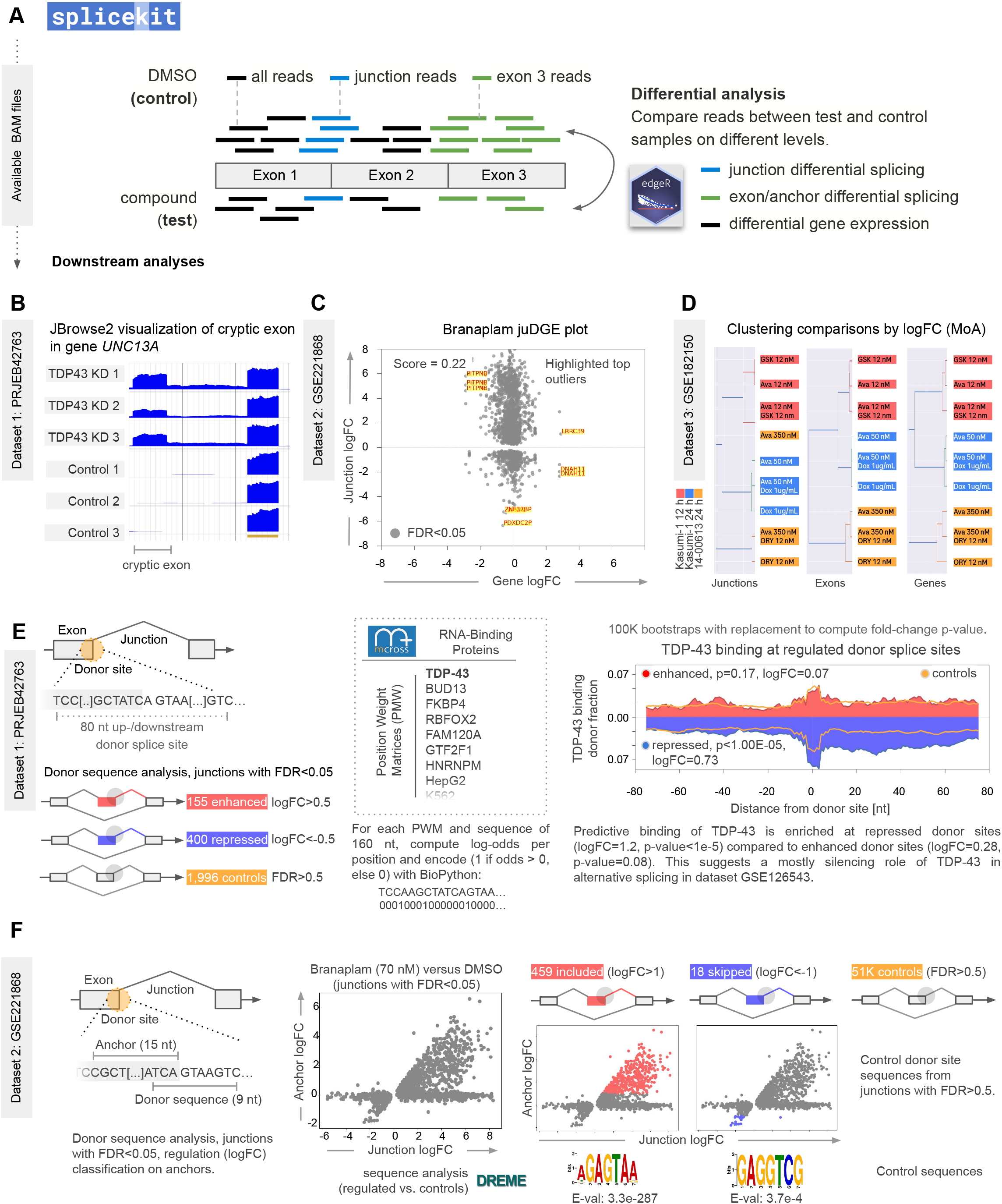
splicekit comprehensive toolkit for splicing analysis from short-read RNA-seq. (A)Initial input to splicekit is read alignments in BAM format. Comparisons are made between groups of test and control experiments. After initial differential calling on the level of splicing (junctions, exons) and genes, further downstream analysis include JBrowse2 visualizations, juDGE plots (logFC of genes and junctions), motif and RNA-protein binding enrichment analysis. (B)JBrowse2 visualization of PRJEB42763 samples and *UNC13A* cryptic exon. The cryptic exon is reported by splicekit junction analysis. (C)junction logFC vs. gene expression logFC (juDGE) plot in GSE221868 suggest Branaplam is a splicing modifying compound. A low juDGE score (stdev_x/stdev_y) suggest most regulation happens at the splicing (junction) level. (D)Clustering of comparisons (test conditions vs. controls) at the junction, exon and gene levels. Only features with FDR<0.05 in at least one comparison. (E)scanRBP analysis of TDP-43 RNA-protein binding in GSE126543 suggests enrichment of TDP-43 binding at repressed sites. (F)Donor Junction Analysis (Don JuAn) on Branaplam versus DMSO in GSE221868 identifies relevant donor sites to detect known binding motif AGAGTAA (Palacino et al., 2015).

## 2 Splicing Analysis

The first step in *splicekit* is to identify regulated features in the comparison, for which *splicekit* runs edgeR (Robinson et al., 2009; Chen et al., 2016) with the diffSpliceDGE function to estimate differential splicing (on junction and exon counts within their respective gene context) and the edgeR glmQLFTest function to estimate differential gene expression.

*splicekit* then integrates diverse existing and novel analysis tools and methods to provide comprehensive differential splicing analysis. It introduces junction-DGE (juDGE) plots, a novel visualization technique to gauge the level of change in the splicing vs. gene context. It implements Donor Junction Analysis (DonJuAn) and motif analysis with DREME (Bailey et al., 2011) to elucidate potential regulatory patterns. In addition, it performs RNA-protein binding scanning (scanRBP) to identify RNA-protein binding in the vicinity of regulated donor and acceptor splice sites.

To visualize read coverage and alignments, *splicekit* provides an integrated JBrowse2 (Diesh et al., 2022) with a containerized web server. Exploring the data in JBrowse2 includes opening the web browser with the local JBrowse2 instance (Figure 1B).

We introduce and further describe novel analysis tools integrated with *splicekit* in sections 2.1 - 2.3.

### 2.1 junction-DGE (juDGE) and cluster logFC plots

To estimate if treated samples display mostly splicing or gene expression changes compared to control samples, *splicekit* produces juDGE plots (Figure 1C). By including genes and junctions and plotting gene log2 fold change (logFC) vs. logFC of junctions, we can estimate the level of alternative splicing in contrast to differential gene expression. A tall vertical plot with a low plot score, defined by the ratio of standard variance of x- and y-axis values, suggests detected changes are mostly on the splicing level, while a wider horizontal plot (high score) suggests there is extensive differential gene expression involved.

In case of multiple comparisons, *splicekit* also provides logFC clustering analysis at the level of junctions, exons and genes. In the Curtiss et al. (2022) dataset, the samples cluster by experimental group (time/cell type) rather than by treatment at all levels (Figure 1D). Such cluster analysis allows an additional overview of the diverse treatments in the context of splicing and gene expression.

### 2.2 Scanning for RNA-protein binding (scanRBP)

To investigate potential involvement of RNA-binding proteins (RBPs) in the mode of action of detected differential splicing events, we established an RBP analysis tool named scanRBP. It plots CLIP data cumulatively (van Nostrand et al., 2016; König et al., 2010) around a set of regulated features or use PWMs (112 RBPs from mCrossAtlas, Feng et al., 2019) to plot log-odds signals for diverse proteins.

The results are RNA-maps that help suggest potential roles of RBPs in splicing changes due to treatment (Rot et al., 2017). Using data reported by Brown et al., 2022, we applied scanRBP to show how TDP-43 represses donor splice site usage (Figure 1E).

### 2.3 Donor Junction Analysis (DonJuAn)

Small molecular splicing modifiers are an arising therapeutic modality that targets specific junction donor sites to modify exon inclusion rates (Sivaramakrishnan et al., 2017). To identify sequence specific splicing effects, we implemented the Donor Junction Analysis (DonJuAn) module in *splicekit*. DonJuAn identifies the donor site sequences and exonic anchor regions of detected junctions for differential expression analysis. Exon inclusion produces positive junction/anchor logFC values, while exon skipping events give negative values which allows for filtering (Figure 1F, left).

As a demonstration, we analyzed public data (Ishigami et al., 2022) on Branaplam, an experimental drug to treat spinal muscular atrophy that binds to donor splice sites, surrounded by the sequence GAGTAAGT (Palacino et al., 2015). DonJuAn logFC junction stratification (Figure 1F, scatterplots) increases the signal for motif enrichment software such as DREME (Figure 1F, DREME).

## 3 Conclusion

We introduce *splicekit*, a comprehensive toolset for analyzing short-read RNA-seq datasets in the context of alternative splicing regulation. By integrating diverse analysis tools and methods, including external tools such as edgeR and DREME, as well as novel tools such as juDGE, DonJuAn, and scanRBP, *splicekit* provides a multifaceted approach to splicing analysis.

One of the key strengths of *splicekit* is its ability to interconnect basic feature analysis with motif search and RNA-protein binding analysis, allowing for a more in-depth understanding of splicing regulation. Additionally, *splicekit* provides exploratory tools for studying the mode of action of splicing in the context of different treatments and compounds.

*splicekit* can be run on a single computer or on a computer cluster, making it a versatile tool for researchers with varying computational resources. Overall, we believe that the scientific community may benefit from adopting, using, and further developing *splicekit*.

